# Oligodendrocyte maturation alters the cell death mechanisms that cause demyelination

**DOI:** 10.1101/2023.09.26.557781

**Authors:** Timothy W. Chapman, Enrique T. Piedra, Robert A. Hill

## Abstract

Myelinating oligodendrocytes die in human disease and early in aging. Despite this, little is known about the mechanisms that govern cell death across the oligodendrocyte lineage. Here we used a combination of intravital imaging, single-cell ablation, and cuprizone intoxication to show that oligodendrocyte maturation dictates the dynamics and mechanisms of death. After single-cell genotoxic damage, oligodendrocyte precursor cells underwent programmed cell death within hours, while mature oligodendrocytes died weeks after the same acute damage. Targeting single cells that were actively undergoing oligodendrocyte generation revealed that a switch in the temporal dynamics and morphological progression of death occurs during differentiation. Consistent with this, cuprizone intoxication initiated a caspase-3-dependent form of rapid cell death in differentiating oligodendrocytes, while mature oligodendrocytes never activated this executioner caspase and exhibited delayed cell death initiation. Thus, oligodendrocyte maturation plays a key role in determining the mechanism of death a cell undergoes in response to the same insult. This means that different strategies are likely necessary to confer protection to the entire oligodendrocyte lineage to enable myelin preservation and facilitate the integration of new oligodendrocytes in aging and disease.

## INTRODUCTION

Oligodendrocytes are susceptible to degeneration in early aging and human disease, resulting in cognitive impairment^1^. Recent work has demonstrated that oligodendrocytes and the myelin sheaths they produce degenerate with surprisingly slow dynamics^2–7^. This has been shown to occur in several demyelination models, as well as spontaneously in the aged brain^8–11^. Despite this, our understanding of the mechanisms that contribute to oligodendrocyte cell death is not well understood.

Several factors contribute to myelin pathology and oligodendrocyte death in aging. First, while oligodendrogenesis from the differentiation of oligodendrocyte precursor cells (OPCs) occurs throughout development and into adulthood, there is an inflection point in oligodendrocyte density, after which newly forming oligodendrocytes fail to properly integrate^8,12–16^. This disrupts the brain’s ability to produce new oligodendrocytes. Simultaneously, established oligodendrocytes begin to degenerate and die. This may be due in part to the accumulation of oxidative damage in oligodendrocytes^17^, but also due to a pro-inflammatory state that microglia and astrocytes take on in response to aging and neurodegeneration^18–20^. These two factors likely initiate a degenerative cascade that exacerbates the neuropathology. Multiple demyelination models are available to study the intrinsic response of oligodendrocytes to external noxious stimuli. Cuprizone intoxication is perhaps the most widely utilized model of demyelination^21^. Despite it having existed for decades, the precise mechanisms that govern oligodendrocyte cell death in the context of cuprizone are not well understood^10,22–24^. This may not be surprising as the exact mechanism by which cuprizone selectively targets oligodendrocytes is also not well understood. And yet, because of its ubiquity in the field, it is critical to understand the mechanisms at play so that accurate conclusions may be drawn, and parallels may be identified with degeneration in aging and disease.

Here we describe how oligodendrocyte maturation shifts the ultimate form of death that individual cells within the oligodendrocyte lineage undergo in response to 2-Photon apoptotic targeted ablation (2Phatal) and cuprizone intoxication. Using 2Phatal, we uncovered a shift in the dynamics of cell death that occurs during the differentiation of OPCs into newly formed and mature oligodendrocytes. We go on to show that a similar effect occurs in cells exposed to cuprizone, suggesting that the observed shift represents an intrinsic change in how oligodendrocytes respond to different noxious stimuli. Finally, we investigated the molecular changes responsible for the differences in death dynamics. Actively differentiating and newly formed oligodendrocytes underwent a caspase3-mediated form of cell death while mature oligodendrocytes died through a distinct, caspase-independent, form of cell death. Previous work has described DNA damage in oligodendrocytes as a characteristic of oligodendrocyte cell death in both cuprizone and human aging. Our data demonstrates that mature oligodendrocytes, and not newly formed oligodendrocytes, do indeed sustain significant DNA damage as was evident from staining for the marker of double-strand breaks, γ-H2AX. Taken together these observations demonstrate a clear switch in the mechanisms of death individual cells undergo as they differentiate and mature. This suggests that distinct therapeutic strategies may be required to protect both the genesis and integration of differentiating oligodendrocytes as well as the previously established mature oligodendrocytes in human aging and disease.

## RESULTS

### Oligodendrocyte maturation alters the temporal dynamics of programmed cell death

2Phatal enables on-demand initiation of programmed cell death in single cells in vivo^25^. We used this precision to target all the cell types across the oligodendrocyte lineage and investigate their cell death dynamics. Before proceeding with this approach, we first needed to precisely identify the maturation stage of the cells under investigation. We used *Cnp-*mEGFP:*Cspg4-*creER:tdTomato mice which label all myelinating oligodendrocytes with mEGFP along with OPCs and their progeny with tdTomato after tamoxifen induced cre recombination (Fig. 1a)^3,13,26^. In addition to labeling OPCs and mature myelinating oligodendrocytes, this mouse line also allows visualization of the cells that are actively differentiating. To identify and define this population, we visualized the differentiation of single OPCs with longitudinal intravital imaging and characterized changes in cell morphology and fluorescence intensities (Fig. 1b). As expected, during active differentiation the *Cnp*-mEGFP signal increased over the course of several days before becoming stable. The cell somas also showed transient and consistent increases in area during differentiation (Fig. 1c). Therefore, plotting soma area as a function of mEGFP fluorescence intensity revealed two oligodendrocyte cell populations which could be separated using unbiased k means clustering (Fig. 1d). In this study we termed these two cell stages as differentiating oligodendrocytes and mature oligodendrocytes. This allowed us to separate our analysis into two oligodendrocyte stages along with OPCs which were identified by their distinct morphology and lack of mEGFP fluorescence.

**Figure 1.**
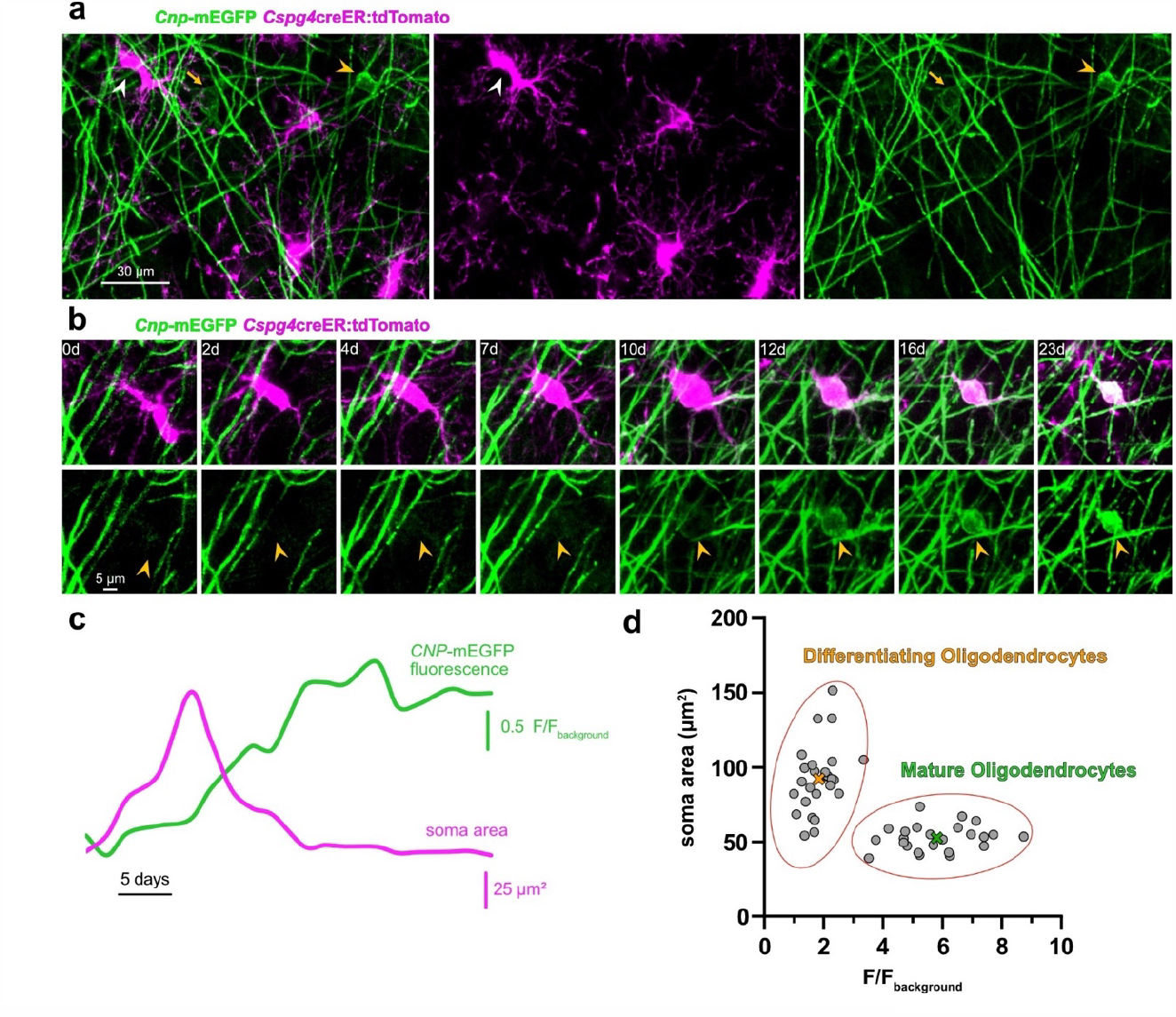
Identifying actively differentiating oligodendrocytes *in vivo*. **(a)** *In-vivo* image of oligodendrocytes and OPCs in layer I somatosensory cortex of a *Cnp*-mEGFP:*Cspg4*creER:tdTomato transgenic mouse. Oligodendrocytes are labeled with membrane tethered EGFP (yellow arrowhead), OPCs are labeled with cytosolic tdTomato (white arrowhead). A newly formed oligodendrocyte is identified by faint EGFP fluorescence and a large soma (yellow arrow). **(b)** Representative timeseries of an OPC differentiating into an oligodendrocyte (arrowhead). EGFP fluorescence signal appears during the differentiation process. **(c)** Trace of normalized EGFP fluorescence over time from the representative timeseries in (b) (green). Trace of soma area of the differentiating cell in (b) over time (magenta). **(d)** Scatter plot of soma area as a function of normalized EGFP fluorescence of oligodendrocytes prior to 2Phatal photobleaching (n=50 cells, 9 animals). Unbiased k-means clustering identified two populations of cells. ‘X’s label the centroids of the two populations, differentiating oligodendrocytes (orange, n=26 cells, 5 mice) and mature oligodendrocytes (green, n=24 cells, 4 mice).

2Phatal works by causing DNA damage in single cells through the photobleaching of nuclear dye in the targeted cell. Hoechst 33342 dye was applied to the cortical surface of the brain during a cranial window surgery in *Cnp-* mEGFP:*Cspg4*creER:tdTomato mice. This enabled us to identify nuclei of OPCs, differentiating oligodendrocytes, and mature oligodendrocytes (Fig. 2). All OPCs targeted underwent an apoptotic-like cell death within 24 hours with nuclear fragmentation and the formation of apoptotic bodies, as previously described^25^. In contrast, mature oligodendrocytes degenerated over an extended period of ∼45 days^3^ without clear morphological characteristics of classical apoptosis, instead exhibiting consistent soma condensation and sheath retraction. On the other hand, differentiating oligodendrocytes demonstrated features and temporal dynamics more similar to OPCs with many dying in 24 hours with others lasting for several days before eventually disappearing (Fig. 2b). There were significant differences across these three groups regarding the temporal dynamics of cell death progression (Fig. 2c, one-way ANOVA, Tukey’s multiple comparison’s test). Thus, the shift in cell death dynamics occurs during differentiation resulting in a population of differentiating oligodendrocytes that display characteristics of OPCs but take longer to die from identical DNA damage protocols.

**Figure 2.**
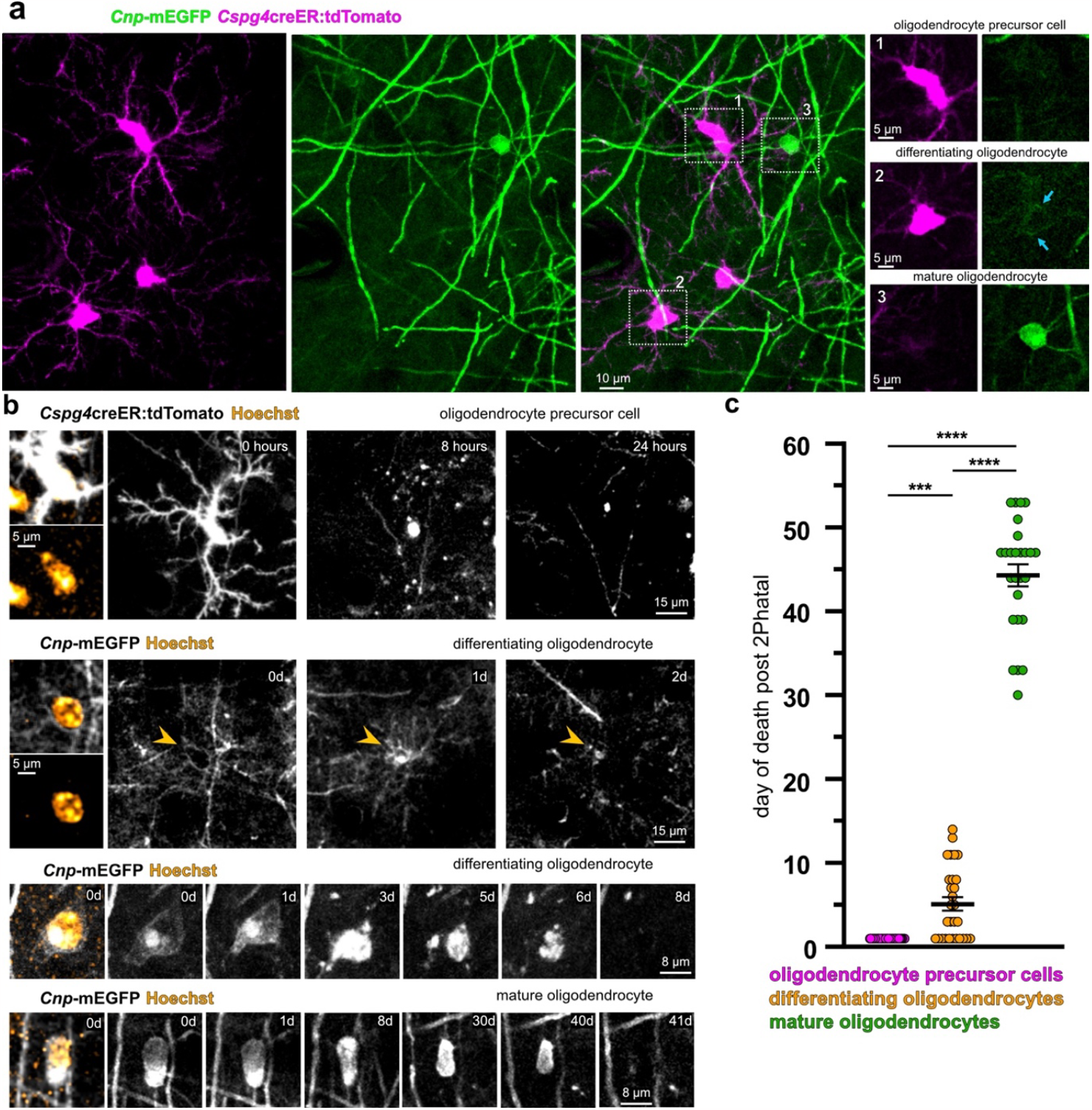
Oligodendrocyte maturation alters the temporal dynamics of induced programmed cell death. **(a)** Representative *in-vivo* image in a *Cnp*-mEGFP:*Cspg4*creER:tdTomato transgenic mouse. Examples of an oligodendrocyte precursor cell (OPC) (box 1), differentiating oligodendrocyte (box 2), and mature oligodendrocyte (box 3) are shown. Cropped boxes are single z sections through the center of the cell soma to indicate the dim mEGFP signal in the differentiating oligodendrocyte (blue arrows) **(b)** Representative timeseries of cells within the oligodendrocyte lineage targeted with 2Phatal. From top to bottom: OPC, differentiating oligodendrocyte (arrowhead identifies the cell soma), differentiating oligodendrocyte, and a mature oligodendrocyte. **(c)** Average day of death post 2Phatal of OPCs (magenta, n=34 cells, 4 mice), differentiating oligodendrocytes (orange, n=26 cells, 4 mice), and mature oligodendrocytes^3^ (green, n=24 cells, 4 mice; one-way ANOVA with Tukey’s correction for multiple comparisons, error bars are SEM)

### Oligodendrocyte maturation defines the cell death dynamics to cuprizone intoxication

We next determined whether oligodendrocyte maturation impacted the degeneration of single cells in the cuprizone model of demyelination^10^. *Cnp*-mEGFP mice were placed on a diet of 0.2% (w/w) cuprizone for 6 weeks and we performed intravital imaging of layer I of the somatosensory cortex. As expected, this regimen produced widespread cortical demyelination after six weeks (Fig. 3a). We identified differentiating and myelinating oligodendrocytes using the criteria described above (Fig. 3c,d, n=40 cells, 3 mice). Mature oligodendrocytes degenerated over an extended period of time, 34 days on average (n=32 cells, 3 mice), as previously described by our lab^3^. Like 2Phatal however, differentiating oligodendrocytes underwent cell death more rapidly than myelinating oligodendrocytes, with an average degeneration time of 14 days (Fig. 3e, n=8 cells, 3 mice, unpaired t-test). We did occasionally observe new oligodendrocyte formation over the course of the cuprizone experiment (Fig. 3b, bottom). These new oligodendrocytes behaved like differentiating cells identified at the start of the experiment, dying more rapidly than previously established mature oligodendrocytes, further indicating that the maturation state dictates the temporal dynamics of cell death between differentiating and mature oligodendrocytes.

**Figure 3.**
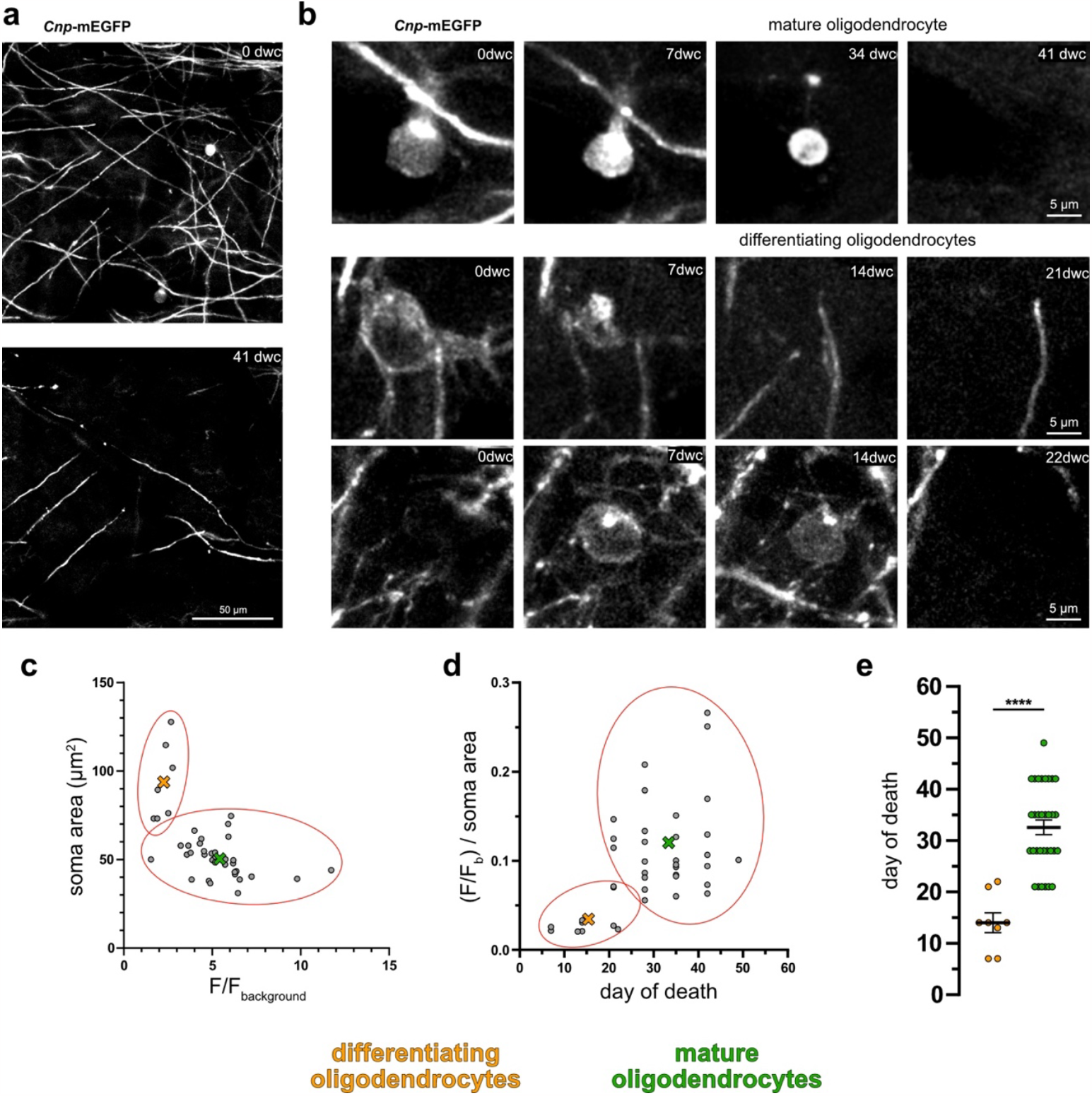
Differentiating but not mature oligodendrocytes die rapidly during cuprizone intoxication. **(a)** *In-vivo* timelapse images showing cuprizone induced demyelination and oligodendrocyte loss in layer I somatosensory cortex over 6 weeks. **(b)** Representative timeseries showing mature oligodendrocyte death (top), differentiating oligodendrocyte death (middle), and the deeath of an oligodendrocyte that appears after the start of cuprizone intoxication (bottom). **(c)** Scatter plot of soma area as a function of normalized EGFP fluorescence of oligodendrocytes prior to the start of cuprizone (n=40 cells, 3 animals). Unbiased k-means clustering identified two populations of cells. ‘X’s label the centroids of the two populations, differentiating oligodendrocytes (orange, n=8 cells, 3 mice) and mature oligodendrocytes (green, n=32 cells, 3 mice). **(d)** Scatter plot made by plotting the ratio of normalized fluorescence to soma area as a function of time to death of each cell, during cuprizone intoxication. Unbiased k-means clustering identified two populations of cells. ‘X’s label the centroids of the two populations. differentiating oligodendrocytes (orange, n=9 cells, 3 mice) and mature oligodendrocytes (green, n=31 cells, 3 mice). **(e)** Average time to cell death of differentiating oligodendrocytes (orange, n=8 cells, 3 mice) and mature oligodendrocytes (green, n=32 cells, 3 mice). The two populations were defined based on the k-means clustering in (c) (unpaired t-test, error bars are SEM).

### Caspase-dependent cell death is limited to differentiating oligodendrocytes

Given the differences in cell death dynamics between differentiating and mature oligodendrocytes revealed with intravital imaging, we next investigated the differences in defined molecular markers of cell death across the oligodendrocyte lineage. To do this we performed another cuprizone experiment in 8-week-old C57bl6/j mice. Mice were placed on a 0.2% (w/w) cuprizone or control diet. Animals were sacrificed after one, three, and five weeks. The brains were then dissected, fixed, and sliced to enable immunofluorescence staining. All imaging for these experiments was done in the somatosensory cortex. It was not possible to use the criteria that defined the oligodendrocyte lineage cell types in our intravital imaging without *Cnp*-mEGFP expression. This meant we had to define the same populations based on molecular markers. We used Platelet derived growth factor receptor alpha (PDGFRα, OPCs), Breast carcinoma amplified sequence 1 (BCAS1, differentiating oligodendrocytes), 2’,3’-cyclic nucleotide 3’ phosphodiesterase (CNP, differentiating and mature oligodendrocytes), and Carbonic anhydrase 2 (CAII, mature oligodendrocytes) to define the specific populations (Fig. 4a,b)^27–29^. Using these markers, we were able to distinguish between differentiating and mature oligodendrocytes, despite both populations expressing CNP (Fig. 4b). We also observed that soma area was larger in the differentiating populations compared to the mature oligodendrocyte population, in agreement with our intravital data (Fig. 4b). However, this characteristic was not used as a defining criterion in our fixed tissue analysis.

**Figure 4.**
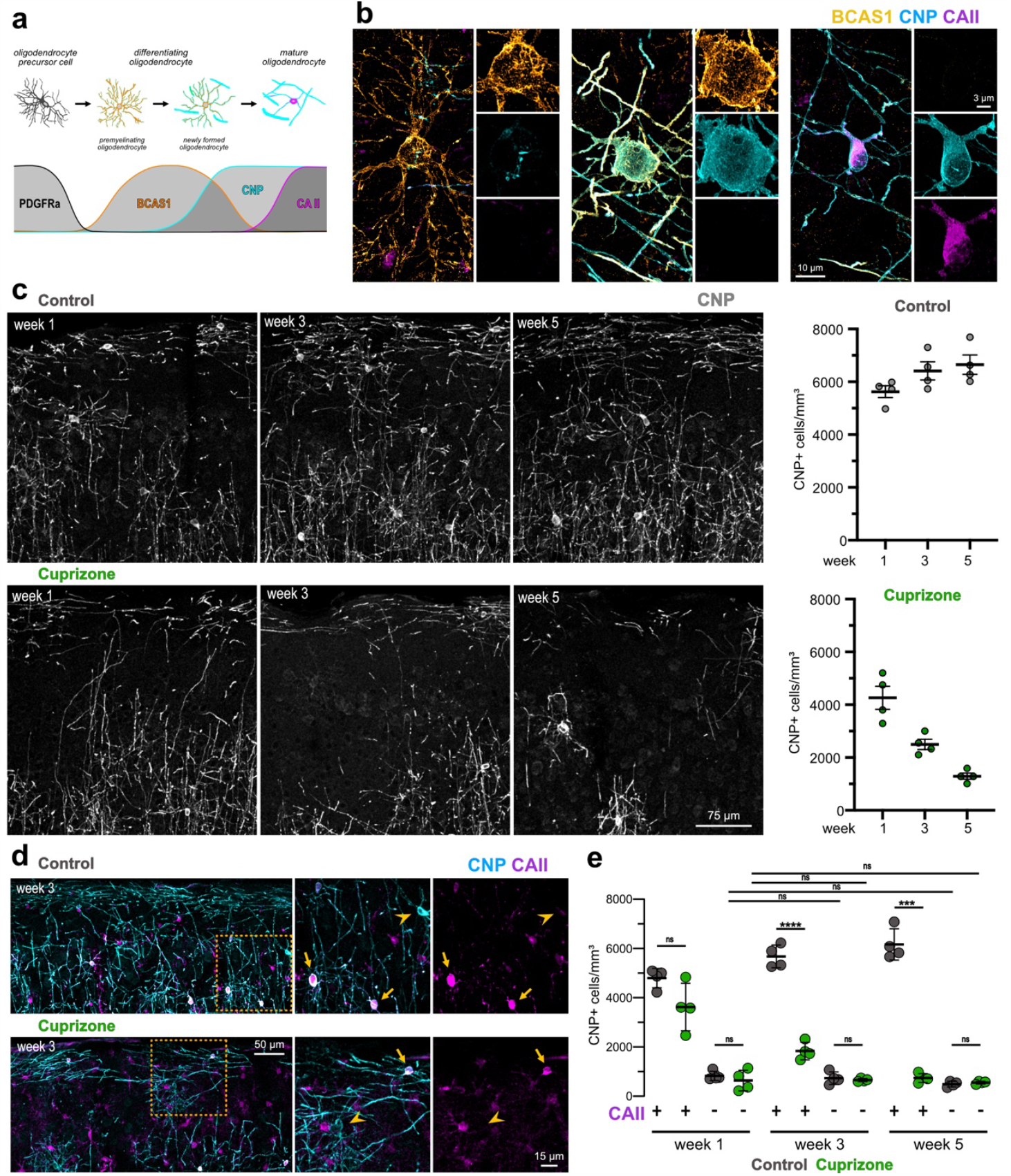
Cuprizone causes failed integration of new oligodendrocytes and delayed death of mature oligodendrocytes. **(a)** Diagram showing the stages of differentiation of the oligodendrocyte linage and the markers used to identify each cell type. Lines represent relative gene expression of each marker in time. **(b)** Representative images of cells within the oligodendrocyte lineage. From left to right: Premyelinating oligodendrocyte (BCAS1+, CNP-, CAII-), newly formed oligodendrocyte (BCAS1+, CNP+, CAII-), and a mature oligodendrocyte (BCAS1-, CNP+, CAII+). **(c)** Representative images and quantification of oligodendrocyte and myelin loss during cuprizone intoxication. Myelin and oligodendrocyte density increase over time in the control condition (n=4 mice), while both myelin and oligodendrocyte density decrease in the cuprizone condition (n=4 mice). **(d)** Representative images of CNP and CAII immunostaining in control (top) and Cuprizone (bottom) tissue from week 3 animals. Zoomed in images show differentiating oligodendrocytes (CNP+CAII-, arrowheads) and mature oligodendrocytes (CNP+CAII+, arrows). **(e)** Quantification of the density of newly formed oligodendrocytes (CNP+CAII-) and mature oligodendrocytes (CNP+CAII+) in control (grey, n=4 mice) and cuprizone (n=4 mice) over time (two-way ANOVA with Tukey’s correction for multiple comparisons, error bars are SEM).

Demyelination was determined by quantifying the density of CNP+ oligodendrocytes in cuprizone and control conditions (Fig. 4c). As expected, there was substantial oligodendrocyte loss and demyelination after five weeks of cuprizone. The oligodendrocyte density of mice in the control condition increased slightly over the 5 weeks, which is consistent with oligodendrogenesis in 8-week-old mice (Fig. 4c). Using the CNP and CAII markers, we next wanted to understand how both newly formed and mature oligodendrocyte populations were contributing to these changes in oligodendrocyte density (Figure 4d). There was no difference in newly formed oligodendrocyte density between control and cuprizone conditions at any time point (Fig. 4e). This suggests that cuprizone does not affect the rate of OPC differentiation into oligodendrocytes. However, unlike the control condition, where differentiating OPCs were able to mature and increase the density of CNP+CAII+ oligodendrocytes over time, newly formed oligodendrocytes in cuprizone were not able to properly integrate contributing to the decline in CNP+CAII+ oligodendrocytes (Fig. 4e).

Previous work has described the appearance of cleaved-caspase3 (CC3) positive cells in the cuprizone model^30^. Interestingly, that work identified two distinct populations of degenerating oligodendrocytes, based on nuclear fragmentation with and without CC3 expression. We sought to understand the role of CC3 in the cuprizone induced cell death progression of cell populations across the oligodendrocyte lineage. Cuprizone and control tissues were stained for CC3, CNP, and CAII (Fig. 5a). Control tissue showed consistently low densities of CC3+ cells at each time point. Tissue from cuprizone treated mice displayed significantly higher densities of CC3+ cells than controls at all three time points. In agreement with previous work^22,30^, CC3+ density was highest after 7 days of cuprizone and sharply fell at weeks three and five, although CC3+ densities continued to be higher in cuprizone than controls at these later time points (Fig. 5c, n=4 mice, one-way ANOVA, Tukey’s multiple comparisons test).

**Figure 5.**
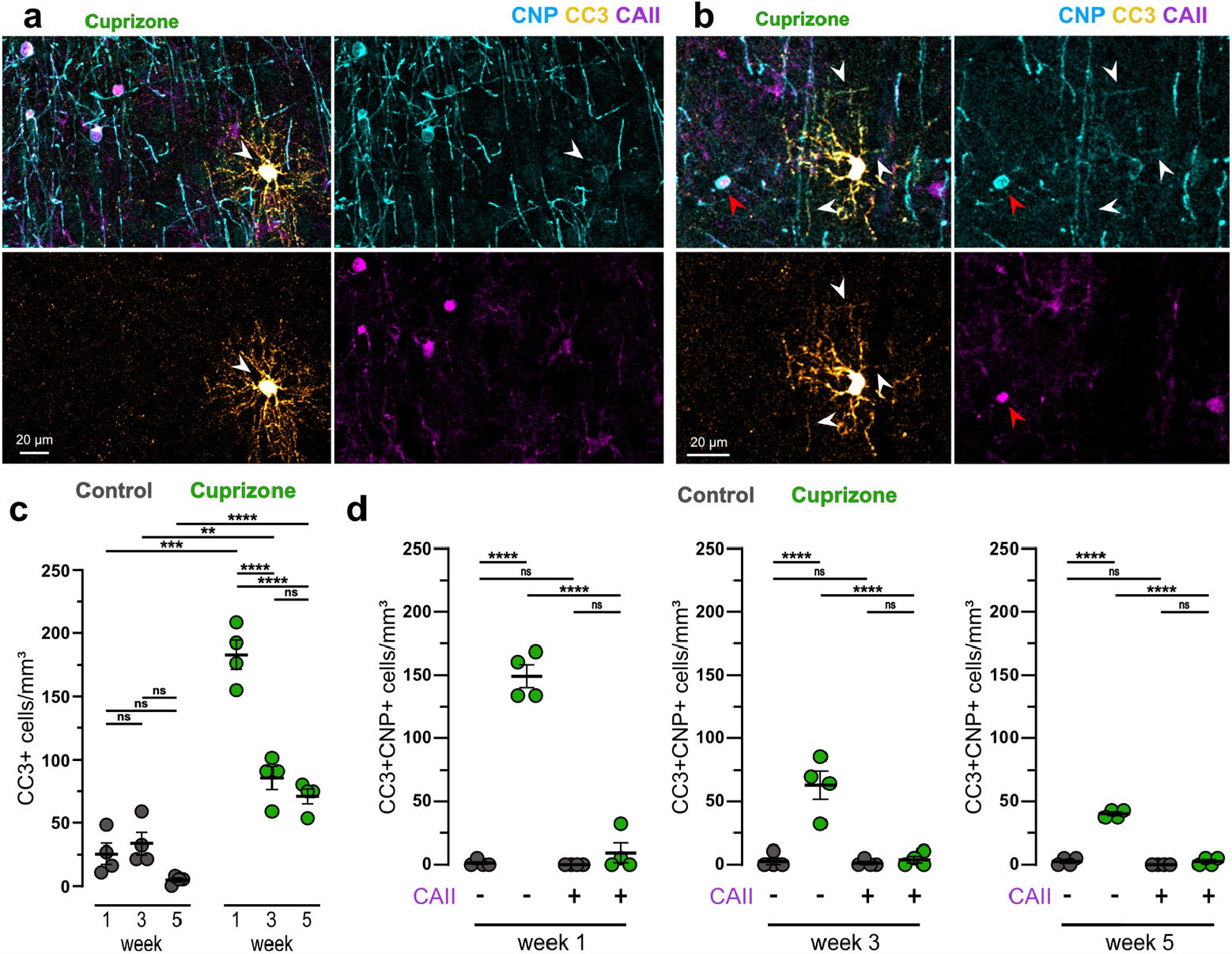
Differentiating but not mature oligodendrocytes die via a caspase3-dependent form of cell death in cuprizone. **(a)** Representative image of a premyelinating oligodendrocyte stained for CNP, CC3, and CAII (arrowhead). The lack of CAII combined with dim CNP signal identifies this cell as a premyelinating oligodendrocyte. **(b)** Representative image of a newly formed oligodendrocyte stained for CNP, CC3, and CAII (white arrowheads). The cell has visible CNP+ myelin sheaths (white arrowheads) but it lacks CAII, identifying this cell as a newly formed oligodendrocyte. **(c)** Quantification of CC3+ cells in the control (grey, n=4 mice) and cuprizone (green, n=4 mice) conditions (one-way ANOVA with Tukey’s correction for multiple comparisons, error bars are SEM). **(d)** Comparison of the density of CC3+ differentiating oligodendrocytes (CNP+CAII-) and CC3+ mature oligodendrocytes (CNP+CAII+) in control (grey, n=4 mice) and cuprizone (green, n=4 mice) conditions. Each graph is a different week of the experiment (one-way ANOVA with Tukey’s correction for multiple comparisons, error bars are SEM).

CC3+ cells were then further separated based on CNP (differentiating and mature oligodendrocytes) and CAII (mature oligodendrocytes) labeling. There was no difference in Caspase3 activation in mature oligodendrocytes between cuprizone treated animals and controls at any time point (Fig. 5d, n=4 mice, one-way ANOVA, Tukey’s multiple comparisons test). In both conditions, CC3+ CNP+ CAII+ densities were either zero or near zero, indicating that mature oligodendrocytes do not contribute to the CC3+ cells observed in cuprizone. In contrast, CNP+ CAII-oligodendrocytes showed increased CC3 activation in cuprizone compared to controls at all time points (Fig. 5d, n=4 mice, one-way ANOVA, Tukey’s multiple comparisons test). These data demonstrate that CC3 activation in cuprizone occurs exclusively in differentiating oligodendrocytes, while mature oligodendrocytes undergo cell death through a caspase3-independent mechanism.

### Cuprizone causes double-strand DNA breaks in mature oligodendrocytes

Oxidative stress and DNA damage in oligodendrocytes have been described in response to cuprizone and aging^31^. We next sought to determine whether the occurrence of DNA damage differed across the oligodendrocyte lineage. The histone protein H2AX has been shown to be phosphorylated at the ser139 residue in response to double strand DNA breaks^32,33^. This modified version of the protein is referred to as γH2AX^34^. As before, we stained tissue from control and cuprizone treated mice for CNP (myelinating oligodendrocytes) and CAII (mature oligodendrocytes), combined with γH2AX (Fig. 6a). As expected, γH2AX signal was confined to the nucleus of positive cells (Fig. 6b). Unlike our previous results with CC3, γH2AX+ cell density was only elevated after 1 week of cuprizone compared to control. There was no significant difference in γH2AX+ cell density between cuprizone and control at weeks 3 and 5 (Fig. 6c, one-way ANOVA, Tukey’s multiple comparisons test).

**Figure 6.**
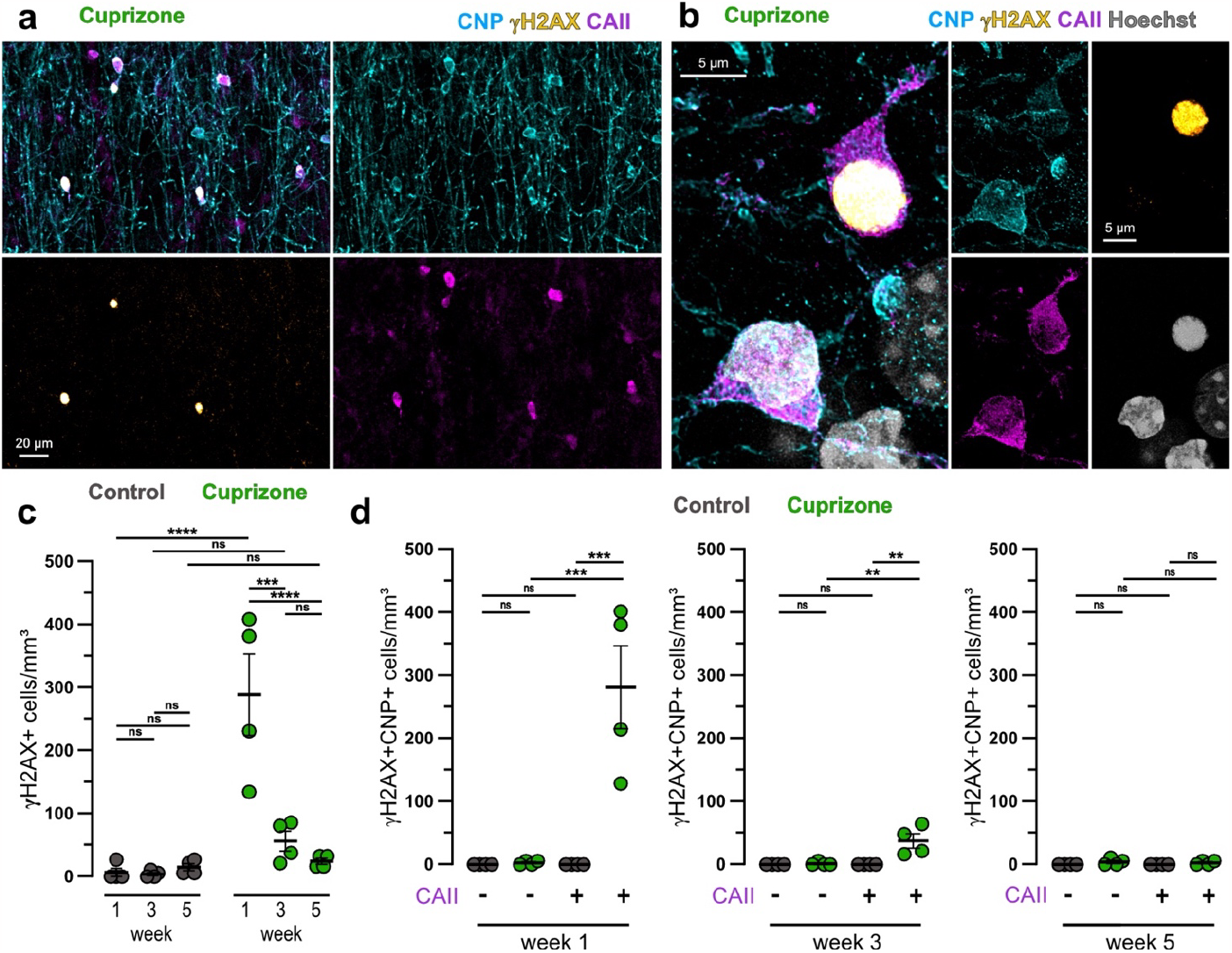
Cuprizone induces double strand DNA breaks in mature but not differentiating oligodendrocytes. **(a**) Representative image of CNP, γH2AX, and CAII staining in a mouse treated with cuprizone. The cells that are positive for γH2AX co-label with CNP and CAII. **(b)** High magnification image of two mature oligodendrocytes. The top cell nucleus is positive for γH2AX while the bottom cell nucleus is not. **(c)** Quantification of γH2AX+ cells in the control (grey, n=4 mice) and cuprizone (green, n=4 mice) conditions (one-way ANOVA with Tukey’s correction for multiple comparisons, error bars are SEM). **(d)** Comparison of the density of γH2AX+ differentiating oligodendrocytes (CNP+CAII-) and γH2AX+ mature oligodendrocytes (CNP+CAII+) in control (grey, n=4 mice) and cuprizone (green, n=4 mice) conditions. Each graph is a different week of the experiment (one-way ANOVA with Tukey’s correction for multiple comparisons, error bars are SEM).

We next separated out the populations of newly formed oligodendrocytes (CNP+ CAII-) and mature oligodendrocytes (CNP+ CAII+) to determine if there was a difference with respect to γH2AX. We did not observe a single γH2AX+CNP+ oligodendrocyte in the control condition at any time point. This is perhaps not surprising as the occurrence of double strand breaks in the healthy brain should be low. However, we also identified only 6 total CNP+CAII-γH2AX+ oligodendrocytes across all three time points in the cuprizone condition (Fig. 6d). The observed γH2AX+ cells in the cuprizone condition were almost exclusively from the mature oligodendrocyte (CNP+CAII+) cell population (Fig. 6d, n=4 mice, one-way ANOVA, Tukey’s multiple comparisons test). These data suggest a transient spike in cuprizone induced DNA damage specifically in mature and not differentiating oligodendrocytes over the first week of cuprizone intoxication which rapidly subsides. Importantly, this occurs well in advance of actual oligodendrocyte degeneration and demyelination (Fig. 4).

### Cortical OPCs do not undergo caspase3-dependent cell death in adulthood or cuprizone

Finally, we determined whether cuprizone had any effect on OPC cell death dynamics. OPCs are generally thought to not be directly affected in the cuprizone intoxication model of demyelination^10^. Many studies have sought to describe the remyelinating response of OPCs during and post oligodendrocyte degeneration in the context of cuprizone. We were interested in understanding whether the switch towards differentiation was required for OPCs to become susceptible to death. To this end, we stained our tissue for PDGFRα (OPCs), BCAS1 (differentiating oligodendrocytes), and CC3 (Fig. 7a). We did not observe any overlap in PDGFRα and BCAS1 labeling indicating that we were not including any differentiating oligodendrocytes in our OPC analysis (Fig. 7b). We were unable to identify any PDGFRa+CC3+ double positive OPCs in tissue from either the cuprizone or control conditions (Fig. 7c,d, n=4 mice, one-way ANOVA, Tukey’s multiple comparison test). This suggests that cuprizone does not directly induce apoptosis in OPCs. The complete lack of CC3+ OPCs in both conditions calls into question whether direct OPC apoptosis plays a substantial role in OPC population homeostasis in the adult cortex.

**Figure 7.**
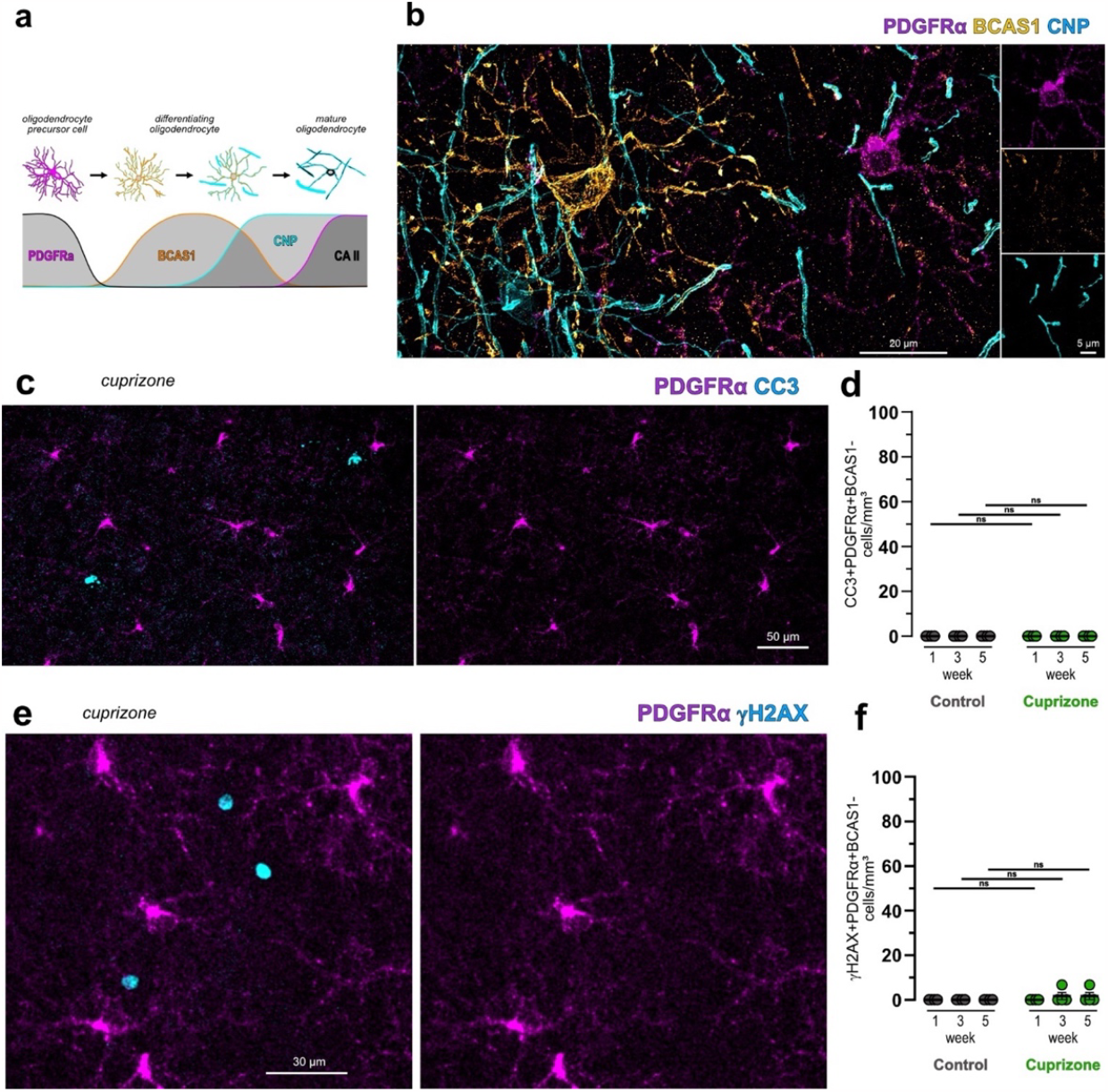
Cortical OPCs do not undergo caspase-dependent cell death in adulthood or with cuprizone. **(a)** Diagram showing the stages of differentiation of the oligodendrocyte linage and the markers used to identify each cell type. Lines represent relative gene expression of each marker in time. **(b)** High magnification image showing an OPC (PDGFRα+, BCAS1-, CNP-), a premyelinating oligodendrocyte (PDGFRα-, BCAS1+, CNP-), and a myelinating oligodendrocyte (PDGFRα-, BCAS1-, CNP+). There is no overlap between PDGFRα and BCAS1. **(c)** Representative image of PDGFRα and CC3 staining in tissue from a cuprizone treated mouse. **(d)** Quantification of the density of CC3+ PDGFRα+ cells in control (grey, n=4 mice) and cuprizone (green, n=4 mice) conditions over time (one-way ANOVA with Tukey’s correction for multiple comparisons, error bars are SEM). **(e)** representative image of PDGFRα and γH2AX staining in tissue from a cuprizone treated mouse. **(f)** Quantification of the density of γH2AX+ PDGFRα+ cells in control (grey, n=4 mice) and cuprizone (green, n=4 mice) conditions over time (one-way ANOVA with Tukey’s correction for multiple comparisons, error bars are SEM).

In addition to CC3, we also investigated whether OPCs sustain DNA damage, like mature oligodendrocytes, as a result of cuprizone intoxication. The previous staining was repeated with γH2AX (Fig. 7e). There was no difference in the density of γH2AX+ OPCs between control and cuprizone conditions (Fig. 7f, n=4 mice, one-way ANOVA, Tukey’s multiple comparisons test) with only a single γH2AX+ OPC was identified at both week three and week five in the cuprizone condition.

## DISCUSSION

When an OPC differentiates into a myelinating oligodendrocyte there are broad morphology changes that are driven by shifts in gene expression, chromatin remodeling, and epigenetic regulation^35–41^. It is therefore not surprising, that cells within this lineage could respond differently to external cytotoxic stress. Indeed, the field has known for decades that cuprizone is selectively toxic to mature oligodendrocytes and not their progenitor cells, and differences across the lineage have been noted in culture and developmental hypoxia-ischemia models^42,43^ . In addition, the fact that the cuprizone intoxication model takes weeks to achieve full demyelination offered a hint into the slow degeneration dynamics of mature oligodendrocytes.

Our goal was to better characterize the cell death of oligodendrocytes as they differentiate and fully integrate, becoming mature oligodendrocytes. First, we showed that the maturation state of individual cells dictates the length of time they take to die in response to 2Phatal induced genotoxic stress. OPCs died within 24 hours while mature oligodendrocytes died 45 days after 2Phatal. Differentiating oligodendrocytes, a category including premyelinating and newly formed oligodendrocytes die with dynamics that fall between these two populations. This demonstrates that the shift in mechanistic response to this kind of insult occurs throughout the differentiating process. We go on to show that the same effect occurs in response to cuprizone intoxication, suggesting that the observed change may represent an intrinsic switch in the overall response to cytotoxic stress. Second, we found that differentiating and mature oligodendrocytes go through molecularly distinct forms of cell death in response to cuprizone intoxication. Differentiating cells use a caspase-dependent form of cell death, which could indicate a more apoptotic like mechanism, while previously integrated oligodendrocytes rarely express cleaved-caspase3. While this finding does not directly identify the precise mechanisms utilized by these two populations, the realization that there is a shift from caspase-dependent to an -independent mechanism clearly identifies a shift in overall mechanism^44^. Finally, we show direct evidence of double strand DNA breaks in mature oligodendrocytes and not in premyelinating and newly formed oligodendrocytes. This again highlights a change in how these populations respond to the same insult. Taken together, these findings highlight specific differences in how cells, within the oligodendrocyte population, respond to cytotoxic stress, and when in maturation that change occurs. It suggests that conferring protection onto these populations, in aging and disease, may require distinct strategies to be maximally effective at preserving previously established oligodendrocytes and enabling the proper integration of new cells.

The accumulation of DNA damage is one of the hallmarks of aging^45^. Oligodendrocytes represent a long lived cell type with a uniquely high metabolic load^14^. This makes them particularly susceptible to age-related stress and spontaneous degeneration. The data we have presented takes advantage of the cuprizone intoxication model and oligodendrocyte 2Phatal to demonstrate distinct differences in the mechanisms by which oligodendrocytes degenerate as a function of maturation. Importantly, while the precise mechanisms that govern cuprizone intoxication are not fully characterized, our data, and that of others, has shown that DNA damage likely plays a substantial role. Indeed, DNA damage is likely the sole driving force in the 2Phatal model. Finally, spontaneous oligodendrocyte and myelin degeneration in aging appears to be a slow process, whereby individual sheaths are lost over time^3,13,46^. This contrasts with what would be expected during a single apoptotic event. It therefore seems likely that oligodendrocytes may degenerate via a caspase-independent mechanism in aging as well. Understanding the precise molecular events that precede this spontaneous degeneration will be critical in the future.

Cuprizone as a model for demyelination has been indispensable for the better part of 60 years. It has been used to great effect, to understand the consequences of demyelination, mechanisms that govern it, and the ability of OPCs to repair it. Despite this, our understanding of precisely how it works remains somewhat of a mystery^10,22^. It is therefore critical to continue to investigate how exactly it induces oligodendrocyte degeneration to accurately draw conclusions from experiments that use it. Here we show that the common oligodendrocyte marker, CNP, captures two separate oligodendrocyte populations that degenerate via distinct mechanisms, during cuprizone intoxication. This finding provides context for several studies that have shown apparently different responses of mature oligodendrocytes to cuprizone^30,47–49^. These studies have produced data that hint at the findings we have presented, including the presence of CC3+ cells in the first weeks of cuprizone^30^. Taken together with our data, these studies produce some intriguing new hypotheses as well. Knocking out the master regulator p53, for example, provides partial protection to oligodendrocytes from cuprizone intoxication^48,49^. One possible explanation for this is that p53 may be required for the death of either the mature or differentiating population of oligodendrocytes. As p53 is known to mediate distinct forms of cell death, if this were to be proven true, it would vastly narrow down the possible pathways involved^50^. In addition, genetic ablation of the BH3 protein, PUMA, has been shown to eliminate the emergence of CC3+ cells in the first 10 days of cuprizone intoxication^47^. This suggests that PUMA may in fact be mediating the caspase-dependent form of cell death that differentiating oligodendrocytes undergo. In line with this hypothesis, knocking out PUMA has been shown to induce hypermyelination in the brain, as well as ectopic myelination in the molecular layer of the cerebellum^51^. As PUMA is known to mediate apoptosis by directly activating BAX/BAK, and thereby triggering caspase activation, it seems logical to propose that the susceptibility of differentiating oligodendrocytes to cuprizone may be due to mechanisms involved in homeostatic cell death. Intriguingly, the same study found that conditional p53 knockout in oligodendrocytes did not induce the same ectopic myelination, further supporting the hypothesis that cell death within the oligodendrocyte linage may occur through both p53-dependent and -independent mechanisms^51^.

The DNA damage response protein, Poly(ADP-ribose) polymerase 1 (PARP1) has also been shown to play a role in oligodendrocyte degeneration. Like the above examples, conditional knockout or molecular inhibition of PARP1 has a protective effect on myelination during cuprizone intoxication^52–54^. However, it does not confer complete protection. It seems likely that this is due to a differential role of PARP1 activation in the death of differentiating and mature oligodendrocytes during cuprizone intoxication.

While we did not observe an increase in CC3+ OPCs at any time during cuprizone intoxication, we also did not observe a single CC3+ PDGFRα+ OPC in the control tissues. This was surprising to us, as caspase3 is generally thought of as the final executioner caspase in the intrinsic apoptosis pathway, and these cells require a mechanism to maintain population homeostasis. These data suggest that population homeostasis in the adult may be achieved almost exclusively as part of the differentiation process instead of at the OPC stage. As differentiation occurs, newly forming oligodendrocytes that are needed, mature and integrate into the myelin network. However, if the new oligodendrocyte is not needed, or there is a negative signal, death occurs, thus achieving OPC population homeostasis without causing ectopic myelination^51,55,56^. This is consistent with the presence of differentiating oligodendrocytes that were CC3+ CNP+ in the control cortex. As our data is limited to neocortical OPCs in the young adult brain, it will be interesting to see if this hypothesis holds true across brain regions and at different developmental stages.

The occurrence of multiple forms of oligodendrocyte cell death during cuprizone intoxication has been previously described in the literature, using differential caspase staining or nuclear morphology^30,57^. This has led to speculation that mature oligodendrocytes may undergo different forms of cell death in response to the same insults^22^. Here, we provide context as to the molecular and cellular mechanisms that govern these observations. Our use of longitudinal imaging of single cells coupled with fixed tissue analysis of defined cell stages revealed that it is the maturation state of oligodendrocytes that dictates the speed and mechanism with which these cells die. As these observations have been made in the context of multiple, distinct, models of oligodendrocyte degeneration, it seems likely that similar mechanisms may govern the death of oligodendrocytes in human aging and disease.

## ACKNOWLEDGEMENTS

We thank members of the Hill lab at Dartmouth for helpful discussions and feedback on the project. This work was supported by the National Institutes of Health R01 NS122800 and the Esther A. & Joseph Klingenstein Fund and Simons Foundation to R.A.H. and a Gilman Fellowship from the Department of Biological Sciences at Dartmouth to T.W.C.

## AUTHOR CONTRIBUTIONS

T.W.C. and R.A.H. conceived, designed, and performed all the experiments and most of the data analysis and quantification. E.T.P. contributed to cell quantification in cuprizone tissue. T.W.C. and R.A.H. wrote the paper and R.A.H. supervised the study.

## METHODS

### Animals

All animal procedures were submitted to and approved by the institutional animal care and use committee (IACUC) at Dartmouth College. The following mouse strains were purchased from Jackson labs and crossed to generate the triple transgenic mice used in this study: *Cnp*-mEGFP^26^ (JAX #026105), *Cspg4*-creER^58^ (JAX #008538), floxed tdTomato Ai9^59^ (JAX #007909). All mouse strains used have C57bL/6 background. Mice were housed in a 12/12 light/dark cycle in a temperature (22 degrees C) and humidity-controlled (30-70% relative humidity) animal vivarium with food and water provided ad libitum. Mice used for fixed tissue cuprizone experiments were 6–8-week-old male, wild-type C57bL/6 mice purchased from Jackson labs. Mice used for 2Phatal and cuprizone intravital experiments were 6-8 weeks old at the start of the experiments.

### Surgical procedures

All intravital imaging was done using chronic cranial window surgical preparations. Briefly, animals were anesthetized with a combination of ketamine (100mg/kg) and xylazine (10mg/kg), given intraperitoneally. The skin over the skull was shaved, sterilized, and removed. A ∼3mm craniotomy was performed using a high-speed drill and the skull was replaced by a #0 cover glass. For 2Phatal, Hoechst 33342 nuclear dye (ThermoFisher #H3570) was diluted to 50 μg/ml in sterile PBS and applied to the pial surface prior to placing the cover glass. A nut was attached to the skull using cyanoacrylate glue to facilitate repeated imaging. The remaining surface of the exposed skull was covered with dental cement. Analgesia was provided by subcutaneous injection of carprofen (50 mg/kg) immediately pre- and post-surgery, and at 24- and 48 hours.

### Imaging

Intravital fluorescence imaging was performed on either a two-photon (Bruker) microscope with an Insight X3 femtosecond pulsed laser (Spectra Physics) using a 20x water immersion objective (Zeiss NA 1.0) or an upright laser scanning confocal (Leica SP8) microscope using a 20x water immersion objective (Leica NA 1.0). We used 488nm to excite mEGFP and 552nm to excite tdTomato. All images with multiple channels were acquired sequentially. For two-photon microscopy, we used 775nm to excite Hoechst dye, 920nm for mEGFP, and 1040nm for tdTomato. Z-stacks were acquired using a step size of 1.5μm. All fixed tissue fluorescence imaging was done on an upright laser scanning confocal (Leica SP8) microscope, equipped with either a 20x air objective (Leica NA 0.75) or a 63x oil immersion objective (Leica NA 1.4). All images with multiple channels were acquired sequentially, from longest to smallest wavelengths. Deconvolution was used on high magnification images (Leica Lightning software).

### 2Phatal

To perform two-photon apoptotic targeted ablations (2Phatal), cranial windows were prepared as previously described. Hoechst 33342 was applied topically (0.1 mg ml^-1^ diluted in PBS) to the pial surface of the brain and allowed to sit for 5 minutes. The cover glass was then secured with cyanoacrylate glue. The windows were allowed to recover for 24 hours to ensure widespread nuclear labeling. Positions were imaged sequentially with 775nm, 920nm, and 1040nm wavelengths and the images merged to enable the identification of nuclei from OPCs and oligodendrocytes. To induce 2Phatal the laser was tuned to 775nm wavelength, with a dwell time of 100 ms, and an ROI (8x8 mm^2^) was centered on the nucleus of interest. Identical time-series parameters were used for each ablation of 125 scans, lasting 3.72 seconds. Following 2Phatal, the positions were reimaged 8 and 24 hours later and then once daily after the first day. Cells to be targeted were identified based on the presence of tdTomato signal and no mEGFP signal for OPCs and for the presence of mEGFP signal for oligodendrocytes. 1 to 9 cells were targeted per position based on the density of labeled cells and the nuclear labeling.

### Cuprizone intoxication

For longitudinal imaging experiments, cranial windows were performed on *Cnp*-mEGFP mice 3 weeks before the start of cuprizone intoxication. Baseline images were acquired on day 0 and then the mice were placed on a 0.2% (w/w) cuprizone (Sigma Aldrich #C9012) diet mixed with ground chow. Food was changed every 2-3 days. Mice were returned to an unadulterated ground chow diet after 6 weeks. For fixed tissue cuprizone experiments, 6-8 week old C57bL/6 male mice were purchased from Jackson labs, allowed to adjust for a week, and then placed on either a 0.2% (w/w) cuprizone diet mixed with ground chow, or an unadulterated ground chow diet. 4 mice from both the control and cuprizone groups were sacrificed once a week for 5 weeks.

### Tissue processing

For tissue dissection, mice were anesthetized with ketamine (100mg/kg) and xylazine (10mg/kg), given intraperitoneally, and transcardiac perfusion was performed with ∼30 ml of 4% PFA in PBS. The brain was then dissected and placed in 4% PFA overnight at 4C, for post-fixation. After ∼24 hours, the tissue was transferred to PBS and stored at 4C. 75μm coronal sections were cut using a vibratome. Tissue was then placed in a glycerol-based cryogenic storage solution at -20C.

### Immunofluorescence staining

Slices were washed 3 times in pbs for 10 minutes to remove cryogenic storage solution. Antigen retrieval was then performed. Slices were placed in a pre-heated solution of 10mM TRIS, 1mM EDTA, and 0.05% TWEEN in PBS buffer at pH 9 for 3 minutes. They were then allowed to cool to room temperature in the solution. The slices were then transferred to primary antibody solution in blocking buffer. Blocking buffer was 0.3% triton-X100 and 1% Bovine Serum Albumin (BSA) and stored at 4C overnight. Slices were washed 3 times in PBS for 10 minutes and transferred to secondary antibody solution in blocking buffer for 1 hour at room temperature. Slices were then washed again for 10 minutes 3 times in PBS. Slices were incubated in a 1:2000 Hoechst solution in 0.3% triton-X100 in PBS for 20 minutes before being mounted using prolong diamond mounting media.

The following primary antibodies were used for immunofluorescence staining: mouse anti-CNPase (Biolegend, 1:500, Cat#836404), Guinea pig anti-CNPase (Synaptic Systems, 1:500, Cat#355004), Rabbit anti-BCAS1 (Synaptic systems, 1:750, Cat#445003), Guinea pig anti-BCS1 (Synaptic Systems, 1:750, Cat#455004), Mouse anti-Carbonic Anhydrase II (Santa Cruz, 1:500, Cat#48351), Goat anti-PDGFRa (R&D, 1:1000, Cat#AF1062), Rabbit anti-Cleaved Caspase3 (Cell Signaling, 1:700, Cat#74001), Rabbit anti-gammaH2AX (Cell Signaling, 1:500, Cat#9718). All staining was done using 0.3% TritonX100 in PBS with 1% Bovine Serum Albumin. Secondary antibodies were conjugated to Alexa Fluor 488, 555, or 647 as necessary (Thermofisher, 1:500).

### Intravital imaging quantification

Changes in soma size of differentiating oligodendrocytes, in *Cnp*-mEGFP:*Cspg4*-creER:tdTomato mice, were quantified using cytoplasmic tdTomato signal. Images were acquired as z-stacks. The slice with the largest soma area was used and the soma was outlined in FIJI. Processes were excluded from the soma area. For the quantification of mEGFP fluorescence, analysis was performed on the same z-slice used for the soma area quantification. The average mEGFP fluorescence of the soma was obtained in FIJI. This was normalized to adjacent background fluorescence intensity in the mEGFP channel. Three background measurements were obtained immediately adjacent to the cell soma. These measurements were averaged, and this average was used to normalize the soma fluorescence value. For quantification of cell death single cells were followed throughout the imaging time series and the day of death was designated as the first day where the cell soma was absent. Raw data for mature oligodendrocytes were reanalyzed from our previous publication^3^ with regard to day of death. New analyses were conducted to determine soma size and mEGFP fluorescence intensity for these cells. All statistical tests were performed in GraphPad Prism or MATLAB and the tests used are indicated in the figure legends and results section.

### Fixed tissue quantification

All fixed tissue analysis was performed using FIJI. Cells were counted and then the volume was calculated to enable the calculation of cell densities. All densities reported were obtained from the cerebral cortex. Experimenter blinding was used for all quantification analyses. For each animal used, three coronal sections were imaged, each 3 times, for a total of nine images per mouse per staining condition. The densities of each of these images were then averaged to produce a single cell density value used for the reported quantification. All statistical tests were performed in GraphPad Prism and the tests used are indicated in the figure legends and results section. Statistical significance is indicated as p values < 0.05.

